# SloR-SRE binding to the *S. mutans mntH* promoter is cooperative

**DOI:** 10.1101/2024.11.02.621577

**Authors:** Myrto Ziogas, India Drummond, Igor Todorovic, Katie Kraczkowsky, Hua Zhang, Hui Wu, Grace Spatafora

## Abstract

*Streptococcus mutans* is a commensal member of the plaque microbiome. It is especially prevalent when dietary sugars are available for *S. mutans* fermentation, generating acid byproducts that lower plaque pH and foster tooth decay. *S. mutans* can survive in the transient conditions of the mouth, in part because it can regulate the uptake of manganese and iron during periods of feast when metal ions are available, and famine when they are limiting. *S. mutans* depends on a 25kDa metalloregulatory protein, called SloR, to modulate uptake of these cations across the bacterial cell surface. When bound to manganese, SloR, binds to palindromic recognition elements in the promoter of the *sloABC* genes that encode the major manganese transporter in *S. mutans*. Reports in the literature describ MntH, an ancillary manganese transporter in *S. mutans*, that is also subject to SloR control. In the present study, we performed expression profiling experiments that reveal coordinate regulation of the *sloABC* and *mntH* genes at the level of transcription. In addition, we describe a role for the *mntH* gene product that is redundant with that of the *sloABC*-encoded metal ion uptake machinery. The results of DNA binding studies support direct SloR binding to the *mntH* promoter region which, like that at the *sloABC* promoter, harbors three palindromic recognition elements to which SloR binds cooperatively to repress downstream transcription. These findings expand our understanding of the SloR metalloregulome and elucidate SloR-DNA binding that is essential for *S. mutans* metal ion homeostasis and fitness in the oral cavity.

## INTRODUCTION

The dental plaque biofilm is home to a diverse microbial community (1, 2) which includes *Streptococcus mutans*, a commensal of the healthy plaque microbiome. Stressors in the human mouth such as a high carbohydrate diet, or a change in oxygen content or pH, can give rise to dysbiotic plaque, where the microbial community is less diverse and associated with disease (3). *S. mutans* is one of the earliest colonizers of the human dentition, and its prevalence on teeth is exacerbated by dietary sucrose which strengthens the interaction of *S. mutans* with the enamel surface (4). Sucrose also serves as a substrate for *S. mutans* homolactic fermentation, the byproducts of which demineralize the tooth enamel and mark the onset of decay (5).

Transition metals such as Mn^2+^ and Fe^2+^ are catalytic cofactors essential for bacterial survival and persistence in a mammalian host (6–8). Metal ion over-accumulation however, can be toxic to cells, thereby necessitating tight regulation of metal ion uptake across the bacterial cell envelope (9). In dysbiotic plaque, *S. mutans* is a uniquely successful pathogen, in part because of its ability to maintain intracellular metal ion homeostasis despite the transient conditions of the human mouth (10). The principal metal ion uptake system in *S. mutans* is the manganese ABC-type transporter encoded by the *sloABC* operon which is modulated by SloR. SloR is a 25kDa transcription factor that is Mn^2+^- dependent and a member of the diphtheria toxin repressor (DtxR) family of metalloregulators. During a mealtime when foodstuffs are plentiful, Mn^2+^ is available to bind to each of three metal ion binding sites in the SloR protein (11). This metal ion sequestration fosters SloR homodimerization and subsequent high affinity binding of SloR to palindromic “SloR Recognition Elements,” or SREs, on the DNA. Specifically, SloR-SRE binding in the *sloABC* promoter region represses downstream *sloABC* gene transcription, thereby preventing intracellular Mn^2+^ over-accumulation that could lead to cell death. During periods of famine when Mn^2+^ is limiting in the mouth, *sloABC* transcription is de- repressed which enables the cells to scavenge this essential micronutrient from the external milieu.

While manganese uptake is primarily mediated by the SloABC metal ion uptake system, Kajfasz et al. (2020) (7) recently described an ancillary transporter of manganese in *S. mutans* that works independent of, yet cooperatively with the SloABC machinery. This ancillary transporter, encoded by the *S. mutans mntH* gene, is an NRAMP-like Mn^2+^-specific permease that, unlike the ABC-type transport that derives from the SloABC system, drives metal ion uptake via secondary active transport. In fact, the *S. mutans* MntH transporter is reminiscent of the NRAMP-like MntH permease in *Bacillus subtilis,* which is transcribed independent of the MntABCD ABC-type system (3). Kajfasz et al. also describe an important role for MntH in optimizing *S. mutans* fitness, and in particular its impact on biofilm formation and oxidative and acid stress tolerance. In that report, *mntH* is described as belonging to the *S. mutans* SloR regulon (7). Specifically, the results of semi-quantitative real-time PCR (qRT-PCR) revealed *mntH* transcription that is 5-fold de-repressed in a *S. mutans* SloR-deficient GMS584 mutant compared to its UA159 wild-type progenitor. Moreover, the results of electrophoretic mobility shift assays (EMSA) support direct SloR binding to the *mntH* promoter region that harbors a predicted SRE immediately upstream of the -10 promoter element (7). Taken together, the SloR-mediated coordinate control of manganese uptake via the SloABC and MntH metal ion transport systems is likely paramount for *S. mutans* survival and persistence in dental plaque and for the cariogenic process that follows. The interaction of SloR with the *S. mutans mntH* promoter region has not been fully characterized however, nor have the mechanism(s) that coordinate MntH- and SloABC- mediated manganese homeostasis been explored.

In the present study, we describe the promoter region of the *S.mutans mntH* gene and the SREs therein to elucidate the details of *mntH* repression by SloR. We propose at least three SREs that localize to the *mntH* promoter region to which SloR binds. We believe this binding is cooperative, involving specific base pair contacts with SloR, as well as protein interactions with the sugar-phosphate backbone. Collectively, the results of this study are consistent with the coordination of redundant SloABC and MntH metal ion transport systems via cooperative SloR binding, and highlight the importance of Mn(II) homeostasis as a means to ensure *S. mutans* survival and persistence in the plaque microbiome.

## MATERIALS AND METHODS

### Bacterial strains, plasmids, and primers

The bacterial strains used in this study are listed in Table 1. Primers and oligonucleotides are described in Table 2 and were designed with Benchling software (2022) using the NCBI *Streptococcus mutans* UA159 genome reference sequence. All primers and oligonucleotides were purchased from Eurofins Genomics (Louisville, KY).

**Table 1.**
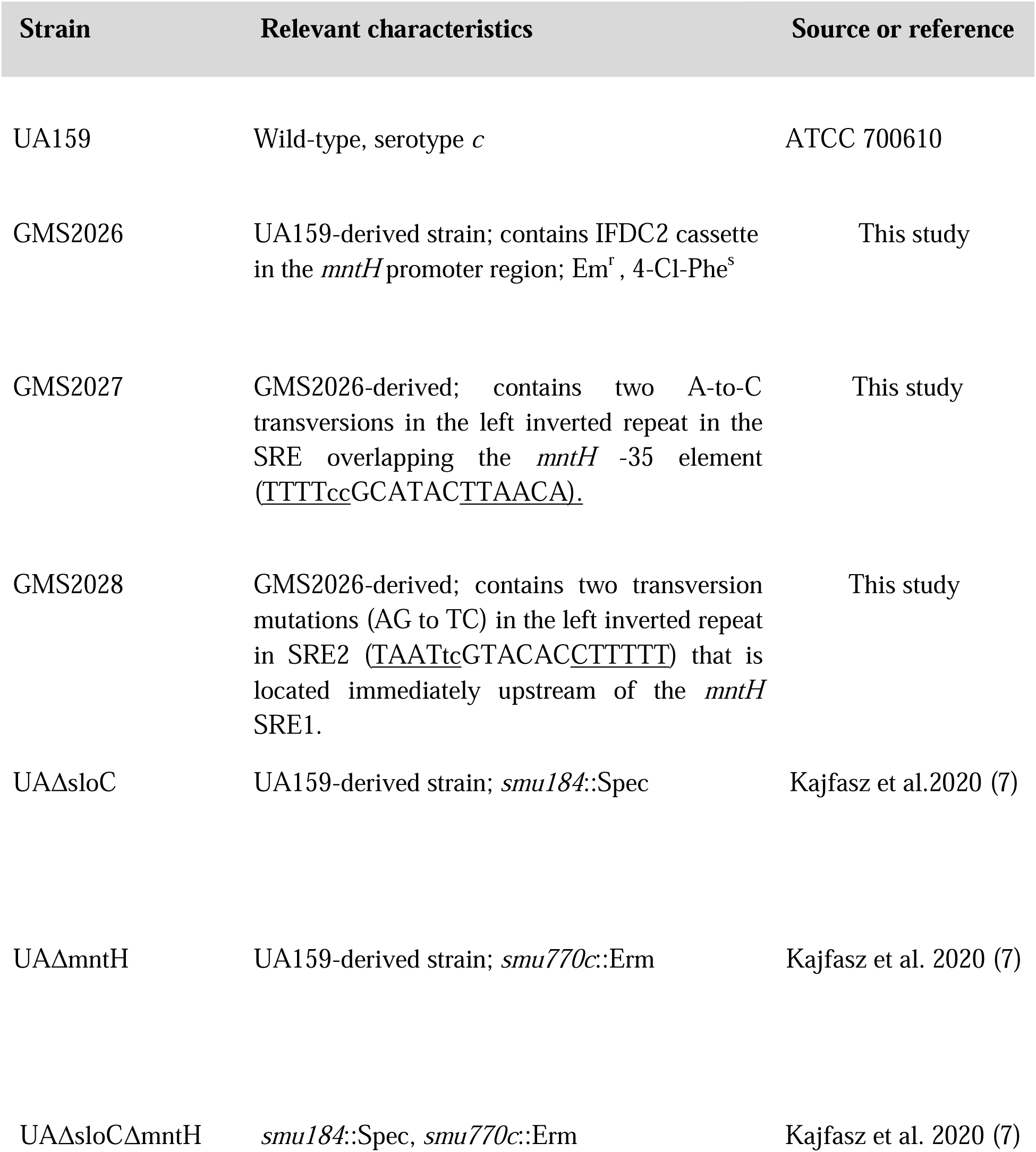
Bacterial strains used in this study.

### Bacterial growth

*S. mutans* was grown as standing cultures at 37°C and 5% CO_2_ in Todd-Hewitt broth (THB) or in THB supplemented with 0.3% yeast extract (THYE). THYE broth was supplemented with erythromycin (10 μg/ml) when growing *S. mutans* GMS2026 which harbors an IFDC2 cassette in the *mntH* promoter region, and with 0.02M p-chloro-phenylalanine (4-CP) when growing SRE variants GMS2027, GMS2028, and GMS2029. All variants in this study were derived from the wild-type *S. mutans* UA159 strain, using the IFDC2 markerless mutagenesis approach described in Xie et al. (12). Briefly, GMS2026 was generated by transforming UA159 cells with the IFDC2 construct and selecting for erythromycin-resistant transformants. Chromosomal DNA was isolated from these transformants and used for PCR and nucleotide sequencing to validate incorporation of the IFDC2 cassette. Transformants were then screened for loss of the IFDC2 cassette on replica plates containing p-chloro-phenylalannine to generate SRE mutant variants GMS2027, GMS2028, and GMS2029 (12). THYE (1/4X) was used to grow *S. mutans* cells for crystal violet biofilm biomass determination assays.

### 54Mn uptake experiments

Standing cultures of the wild type *S. mutans* UA159 strain, and its Δ*sloC*, Δ*mntH,* and Δ*sloC/*Δ*mntH* derivatives were grown in 14 mL of Todd Hewitt broth (THB) and incubated at 37°C with 5% CO_2_ overnight. 250 μl of each culture was subsequently transferred to separate sterile Falcon tubes containing 45 mL of pre-warmed THB, and incubated until early exponential growth phase (OD_600nm_ = 0.1). All culture densities were normalized to within 0.005 OD units before transferring 1 mL of each culture to 1.5 mL microcentrifuge tubes and adding 1 μL of 1.8 μCi/μL ^54^Mn (or 1 μL of 0.5 M HCl for the control). All cultures were incubated as described above for 16-18 hours, after which the cells were pelleted by centrifugation and the supernatants for each experimental sample were set aside for scintillation counting. The cell pellets in both the experimental and control groups were sequentially washed in 1 mL of THB prior to resuspension. Variation in the experimental design includes some inevitable loss of cell- associated ^54^Mn during these wash steps. 100 μL of each cell suspension was transferred to vials containing scintillation fluid and assessed for counts per minute (cpm) in a liquid scintillation counter. Counts per minute for 100uL samples of each supernatant were determined in parallel. For the control sample, 10-fold serial dilutions were plated onto THB agar plates and viable plate counting was performed to determine the number of colony forming units (cfu) in each sample.

### Nucleic Acid Isolation

Chromosomal DNA and total intact RNA were isolated from *S. mutans* cells according to established protocols (13, 14).

### 5’ RACE

To identify the *mntH* transcription start site and predict the locale of the -10 and -35 promoter elements that drive *mntH* transcription, we performed 5’ Rapid Amplification of cDNA Ends (5’ RACE) using total RNA from *S. mutans* UA159 and an Invitrogen 5’ RACE kit (15). The primers used in this study are shown in Table 2 and include 770.GSP1.2 for reverse transcription, 770.GSP2.2 for the first round of PCR amplification, and 770.GSP3.2 for the nested PCR amplification. The reaction conditions for both PCR rounds included Platinum HiFi Taq Polymerase and a 94°C hot start, followed by a 1 minute incubation at 94°C and then 35 cycles of 94°C for 15s, Ta for 30s (55°C for the first round of amplification and 54°C for the second round), 68°C for 45s, and a 4°C hold. Samples were outsourced for sequencing (Eurofins, Inc.) and results were aligned with the UA159 reference genome (RefSeq accession number NC_004350.2) using Benchling to define the transcription start site.

### DNase I footprinting

DNase I footprinting was performed according to established protocols (Spatafora et al. 2015). A 291-bp amplicon containing the *mntH* promoter region was PCR-amplified with primers mntH.F1.fp and mntH.R1.fp. The former was end-labeled with 32P-ATP and T4 polynucleotide kinase (Table S1) and the resulting amplicon was purified on a Qiaquick PCR column according to the manufacturer’s instructions (Qiagen), and stored at -20°C in nuclease-free H_2_O. Binding reactions were performed by combining 10 uL of 5X binding buffer (final concentration 8.4 mM NaH_2_PO_4_, 11.6 mM Na_2_HPO_4_, 50 mM NaCl, 5 mM MgCl_2_, 10 ug/mL bovine serum albumin, 200 ug/mL salmon sperm DNA, and 7.5 uM MnCl_2_) and up to 2 uM purified native SloR protein (Spatafora et al. 2015) in a 45 uL total volume. The protein and its binding buffer were incubated at room temperature for 10 minutes, after which 5 uL of radiolabeled amplicon was added. The reaction mixtures were incubated at 30°C for 30 minutes after which 5 uL of RQ1 DNase I (0.02 Kunitz units/uL) and 6 uL of 10X RQ1 reaction buffer were added for subsequent incubation at 37°C for 45 seconds. Stop solution (20 mM EGTA [pH 8.0]) prewarmed to 37°C was then added and the reaction mixture incubated for 10 minutes at 65°C. The DNA was purified via phenol chloroform extraction and precipitated for 5 days at -20°C in 400 uL of 100% ethanol. The DNA-protein complex was pelleted by centrifugation at 14,500 rpm for 30 minutes and 4°C and the pellets were washed in 70% ice-cold ethanol and dried for 10 minutes in a vacufuge at 30°C. The pellets were resuspended in Stop/Loading buffer (0.1% [wt/vol] bromophenol blue, 0.1% [wt/vol] xylene cyanol, 10 mM EDTA, 95% [vol/vol] formamide) before loading (2 uL per lane) on an 8% urea-containing polyacrylamide gel alongside the sequencing reactions (2.5 uL per lane). Sequencing reactions were performed with an Affymetrix Thermo Sequenase Cycle Sequencing Kit in accordance with the manufacturer’s instructions for radio-labeled primer cycle sequencing (USB, Cleveland, OH). The DNA footprint was resolved for 1.5 hours at 1,600 V, after which the gel was loaded a second time and run for an additional 1.5 hours. The gel was exposed to Kodak BioMax film for up to 24 hrs at -80°C in the presence of an intensifying screen before it was developed by autoradiography.

### Electrophoretic Mobility Shift Assays (EMSAs)

EMSAs were performed using established protocols (Haswell et al. 2013) to define the region in the *mntH* promoter to which SloR binds. Primers or oligonucleotides were designed to generate individual target probes that collectively span the 206-bp *mntH* promoter region or derivatives thereof. PCR reactions with Q5 High-Fidelity DNA polymerase (New England BioLabs) were used to amplify the target probes from the *S. mutans* UA159 chromosome. Amplicons were confirmed by agarose gel electrophoresis and purified with a Qiagen PCR Purification Kit (Thermo Fisher, Waltham, MA) prior to end-labeling with γ-32P-ATP in the presence of T4 polynucleotide kinase (New England BioLabs). Single-stranded oligonucleotides were annealed according to established protocols before end-labeling (16). Binding reactions were prepared as described previously (15) in a 16ul reaction volume containing purified native SloR protein (10μg/ml) at concentrations ranging from 60nM to 200nM. EDTA was added to select reaction mixtures at a final concentration of 15 mM to confirm that SloR-DNA binding was Mn-dependent. A reaction mixture containing a target probe on which the *sloABC* promoter is resident was used as a positive control for SloR binding (7, 17). The samples were loaded onto 12% nondenaturing polyacrylamide gels and run for 450 to 600 Volt-hours depending on the length of the DNA probe.

Gels were exposed to Kodak BioMax film for up to 5 days at -80°C in the presence of an intensifying screen for subsequent autoradiography.

### Construction of *S. mutans mntH* SRE variant strains

To introduce site specific mutations into the *mntH* promoter region, we used *S. mutans* UA159 and a markerless mutagenesis approach described previously by Xie et al (12). Specifically, we performed overlap extension PCR (OE-PCR) to generate a linear construct which contained a 2.2 kb IFDC2 cassette with homologous arms that flank the *mntH* promoter region. This construct was transformed into *S. mutans* UA159 with competence-stimulating peptide (CSP) and integrated into the chromosome via allelic exchange to generate GMS2026, an erythromycin-resistant and *p*-4-chlorophenylalanine (*p*-Cl-Phe) -sensitive derivative. The double-crossover event was confirmed by PCR and Sanger sequencing. A derivative of GMS2026 was subsequently generated by introducing two point mutations into *mntH* SRE1 followed by allelic exchange into the GMS2026 chromosome (Table 2). The mutated OE-PCR construct was prepared with a degenerate primer set harboring A→C mutations that localize to the leftmost inverted repeat of SRE1 upstream of the *mntH* -35 promoter. Incorporation of this mutation into the *S. mutans* chromosome at the locale of “SRE1” was validated by Sanger sequencing, and the resulting strain was called GMS2027. In a parallel set of experiments, two other GMS2026 derivatives were generated as OE-PCR constructs containing two transversion mutations in the leftmost hexameric repeat of so called “SRE2” (<u>TAATAG</u>GTACAC<u>CTTTTT</u> to <u>TAATtc</u>GTACAC<u>CTTTTT</u>) or mutations in both SRE1 and SRE2. The resulting mutant variants were named GMS2028 and GMS2029, respectively.

### Semiquantitative real-time PCR (qRT-PCR)

To assess the impact of SRE mutations on downstream *mntH* transcription, we isolated total intact RNA from *S. mutans* UA159, GMS2027, GMS2028, and GMS2029 cultures grown to mid-logarithmic phase in THYE (the UA159-derived mutant variants were grown in a medium supplemented with *p*-4-chlorophenylalanine). RNA quality was assessed on an Agilent Bioanalyzer before reverse transcribing 100 ng of each RNA into cDNA. The cDNAs were used as templates in qRT-PCR experiments performed in a CXR thermal cycler (Bio-Rad) according to established protocols (16) The expression of *mntH* was measured in each of three independent experiments, each performed in triplicate and normalized against the expression of an *hk11* gene, which did not change under the experimental test conditions (18).

### Phenotypic characterization of *S. mutans mntH* mutant strains

#### Crystal Violet Release Assays

Overnight cultures of *S. mutans* UA159, its isogenic *mntH*, *sloC*, and *mntH/sloC* insertion deletion mutants were all grown to mid-logarithmic phase (OD_600nm_ of 0.4–0.6) in THYE medium at 37°C and 5% CO_2_. The cells were centrifuged at 7000 rpm for 4 min and the resulting pellets resuspended in 10mL of 1/4 strength THYE containing 18mM glucose and 2mM sucrose. The resuspended cultures were used to inoculate a 24-well polystyrene microtiter plate (Corning, New York, USA) containing fresh 1/4X THYE medium as described above. The microtiter plates were incubated for 18 hrs to allow for biofilm formation, and then agitated gently for 5 minutes on a Gyrotory Shaker-Model D2 rotating platform (New Brunswick Scientific Company, NJ, USA) set at 100 rpm to dislodge non-adherent cells. Culture supernatants were decanted by gentle inversion and the biofilms were washed once with 2 mL of water. 500µL of crystal violet (0.1% w/v) was then added to each well and the plates were incubated standing at room temperature for 15 min. Excess crystal violet was removed from the wells by gentle inversion, and the wells were washed twice with 1 mL of water before they were air-dried for 1hr. Intracellular crystal violet was extracted from the adherent cells in 2 mL of 99% ethanol and absorbance was measured at 575 nm in a Genesys 20 spectrophotometer (Thermo Scientific). Wells containing uninoculated 1/4X THYE were included in the experimental design and used as negative controls.

#### Growth Determination Assays

Overnight cultures of *S. mutans* UA159 and its mutant derivatives were grown to mid-logarithmic phase (OD_600nm_ 0.4–0.6). Final cell culture densities were standardized to within 0.05 OD units with fresh THYE broth before inoculating microtiter wells (1:100) in duplicate in THYE medium. Growth was monitored at 600 nm over a 24 hour period in a BioScreen C Microplate Reader (Thermo Labsystems) according to the manufacturer’s instructions.

#### Scanning Electron Microscopy (SEM)

Overnight cultures of *S. mutans* UA159 and its mutant derivatives were grown to mid-logarithmic phase (OD_600nm_ 0.4–0.6) in THYE medium as described above. Biofilm growth was initiated by inoculating mid-logarithmic phase cells into the wells of a 24- well polystyrene microtiter plate, each containing fresh THYE medium and sterile Thermanox coverslips (Electron Microscopy Sciences). The biofilms were grown for 18 hrs at 37°C and 5% CO_2_, washed once in 10mM phosphate-buffered saline (PBS), fixed at room temperature in 2mL of 3.7% formaldehyde in 10mM PBS for 24 hours, and then dehydrated in ethanol rinses prior to air drying. All samples were sputter coated with a gold-palladium mixture and then examined with a Tescan Vega 3 LMU scanning electron microscope (Brno, Czech Republic).

#### Biolayer Interferometry

Biolayer Interferometry (BLI) was performed using the Octet Red96 system (FortéBio, Fremont, CA) as previously described (19). In brief, kinetic binding assays were performed at 30°C with a shaking speed of 1000 rpm in a 384-well plate. Each well contained 60 µL of phosphate-buffered saline (PBS, pH 7.2) supplemented with 0.1% Tween-20 and 100 µM MnCl_2_.

Purified SloR protein in phosphate-buffered saline (PBS, pH 7.2) was immobilized on an NTA biosensor, followed by the application of a series of diluted synthesized oligonucleotide probes through the protein-coated biosensors. Binding interactions between SloR and the probes were analyzed using the Octet Red System Data Analysis software.

## RESULTS

### The *S. mutans* MntH and SloABC manganese transport systems are redundant and compensatory

To elucidate the relative contribution of MntH-mediated Mn^2+^ transport to total intracellular manganese in *S. mutans*, we conducted ^54^Mn uptake assays with the UA159 wild-type strain and its derivatives harboring mutations in genes that encode metal ion transport. Specifically, these assays monitored relative intracellular ^54^Mn concentration across strains by comparing the counts per minute associated with the *S. mutans* UA159 cell pellet with that of its Δ*mntH*, Δ*sloC,* and Δ*mntH*Δ*sloC* derivatives. The experimental findings reveal heightened accumulation of ^54^Mn in the Δ*mntH* cell pellet compared to wild type, and even greater ^54^Mn accumulation in cell pellets of the double mutant (Fig. 1). The latter is consistent with the presence of additional inducible metal ion uptake systems in *S. mutans*, and the former with a compensatory role for the SloABC- and MntH-mediated manganese transport systems when either *mntH* or *sloABC* is compromised. Compensatory ^54^Mn uptake in the Δ*mntH* mutant is significantly greater than that of the Δ*sloC* mutant, consistent with an ancillary role for manganese uptake via the MntH transport mechanism.

**FIG 1.**
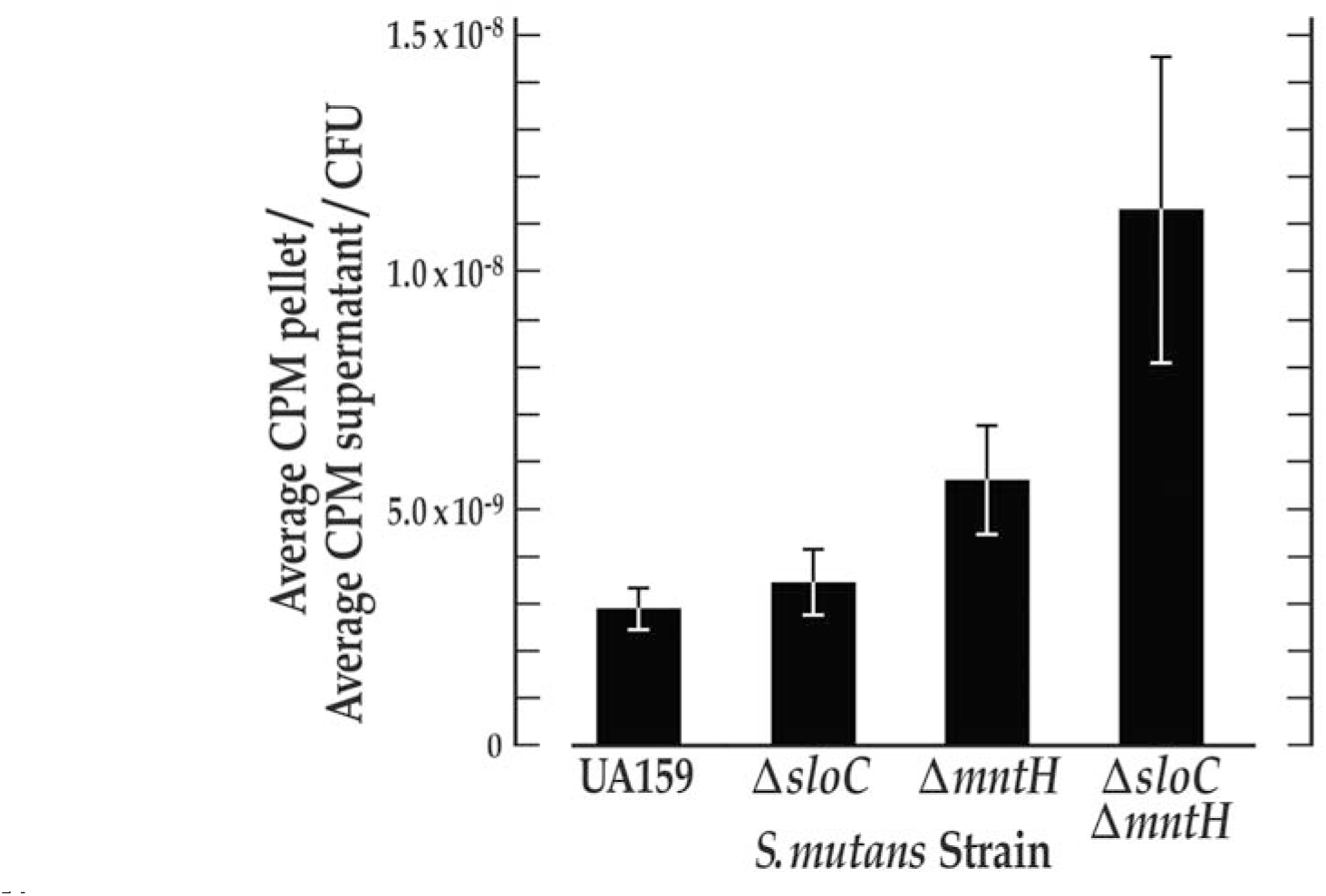
^54^Mn uptake experiments support SloC and MntH as compensatory transporters of manganese in *S. mutans*. ^54^Mn accumulates most notably in the *S. mutans* Δ*sloC*/Δ*mntH* double mutant when compared with that of its UA159 wild-type progenitor or its Δ*sloC* or Δ*mntH* single mutant derivatives. All *S. mutans* cultures were grown overnight to within 0.005 absorbance units in the presence of ^54^Mn or a 0.5M HCl control. The cell pellets and supernatants were separated by centrifugation and counts per minute (CPM) were determined by liquid scintillation counting. The number of colony forming units (CFU) was determined by plating serial dilutions of the experimental and control cultures in parallel. The CPM of the cell pellets were divided by the CPM of the supernatants to correct for cell loss, and that ratio was then divided by the number of CFUs to normalize the ^54^Mn transport output. Error bars denote the standard deviation about the mean.

To determine whether compensatory roles for the *mntH* and *sloABC* systems could be demonstrated at the level of phenotype, crystal violet release assays were performed to assess the biofilm biomass of the *S. mutans* wild-type and mutant strains. These experiments were performed in parallel with scanning electron microscopy imaging to monitor the details of biofilm architecture in these strains. Growth determination assays were also performed to ensure similar rates of growth for *S. mutans* UA159, and its isogenic Δ*mntH,* Δ*sloC,* and Δ*mntH*Δ*sloC* manganese uptake mutants. The results of growth determination assays confirmed similar doubling times (90 minutes) for all test strains (data not shown). Crystal violet assays revealed a significant reduction in biofilm biomass for the *mntH*/*sloABC* double mutant compared to its UA159 wild-type progenitor and single mutant derivatives (Fig. 2a and b). In contrast, the biofilm biomass of the Δ*mntH* and Δ*sloC* single mutants was not significantly different from that of the wild-type, consistent with the compensatory roles of the MntH and SloABC Mn2+ uptake systems. Interestingly, scanning electron micrographs of the *S. mutans* UA159 and Δ*mntH*Δ*sloC* strains revealed very different biofilm architectures (Fig. 2c). Specifically, the Δ*mntH*Δ*sloC* mutant was nearly devoid of water channels, in stark contrast to the many water channels that were observed in the UA159 strain. The biofilm architecture of the Δ*mntH* and Δ*sloC* single mutants was indistinguishable from that of UA159 wild-type (data not shown), lending further support to compensatory roles for the MntH and SloABC systems in *S. mutans*.

**FIG 2.**
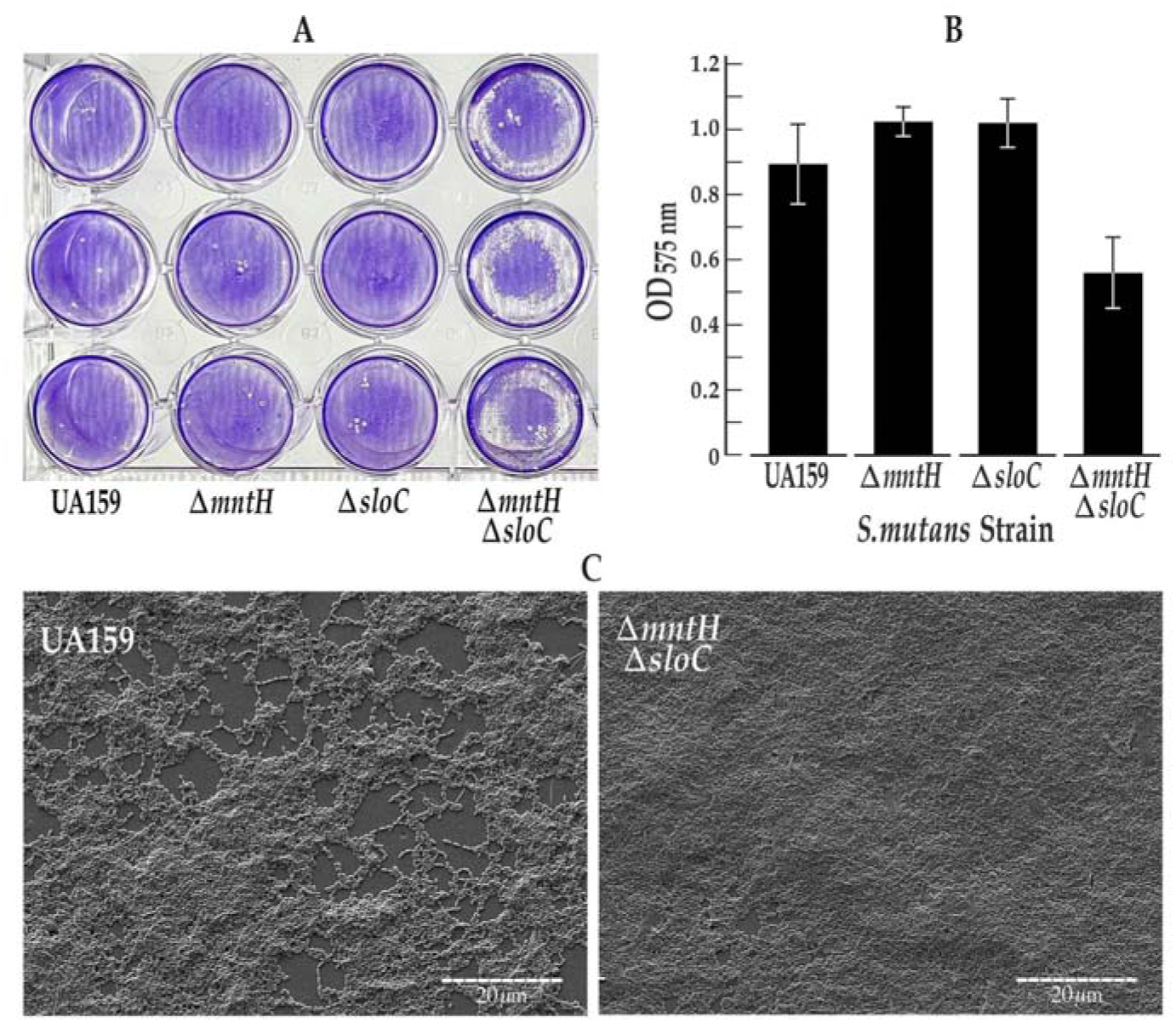
The *S. mutans* Δ*mntH*Δ*sloC* mutant is compromised for biofilm formation. Cultures were grown overnight to mid-logarithmic phase in 1:4 THYE media supplemented with 18mM glucose and 2mM sucrose. (a) Results of crystal violet assay demonstrate reduced biofilm formation in the Δ*mntH*Δ*sloC* mutant compared to the UA159 wild-type and its single mutant derivatives (b) Spectroscopy of crystal violet extracts from adherent biofilm cells reveals significantly compromised biofilm biomass for the Δ*mntH*Δ*sloC* mutant compared to the UA159 wild-type strain (p=0.02), and its isogenic Δ*sloC* (p=0.004), and Δ*mntH* mutants (p=0.002). Error bars depict standard error of the mean. (c) Scanning electron micrographs of *S. mutans* UA159 and Δ*mntH*Δ*sloC* biofilms grown on polystyrene coverslips reveal marked differences in biofilm architecture. Most notably, the Δ*mntH*Δ*sloC* biofilms lack the water channels that are present in the UA159 wild-type biofilms. Shown are representative regions of the electron micrographs for *S. mutans* UA159 and its isogenic Δ*mntH*Δ*sloC* mutant. Biofilm architecture of the single mutants was indistinguishable from that of the UA159 wild-type progenitor (data not shown).

### Characterization of the *S. mutans mntH* promoter region

To inform gene regulation at the *mntH* locus we characterized the *mntH* promoter region by defining the transcription start site (TSS) from which we could predict SRE positioning relative to the - 10 and -35 promoter elements. To this end, we conducted 5’ RACE experiments, the results of which support *mntH* transcription that begins at an adenosine residue located 32 bp upstream of the ATG start codon (Fig. 3). Based on the location of this adenosine residue, we predicted the positioning of the -10 and -35 *mntH* promoter elements, while taking into consideration the -10 and -35 consensus sequences in prokaryotes (20). The predicted -10 promoter element has a sequence that exactly matches that of the canonical prokaryotic sequence (TATAAT), while the predicted element positioned 35 nucleotides upstream of the TSS (TTAACA) deviates from the conserved -35 prokaryotic sequence (TTGACA), but only by a single nucleotide.

**FIG 3.**
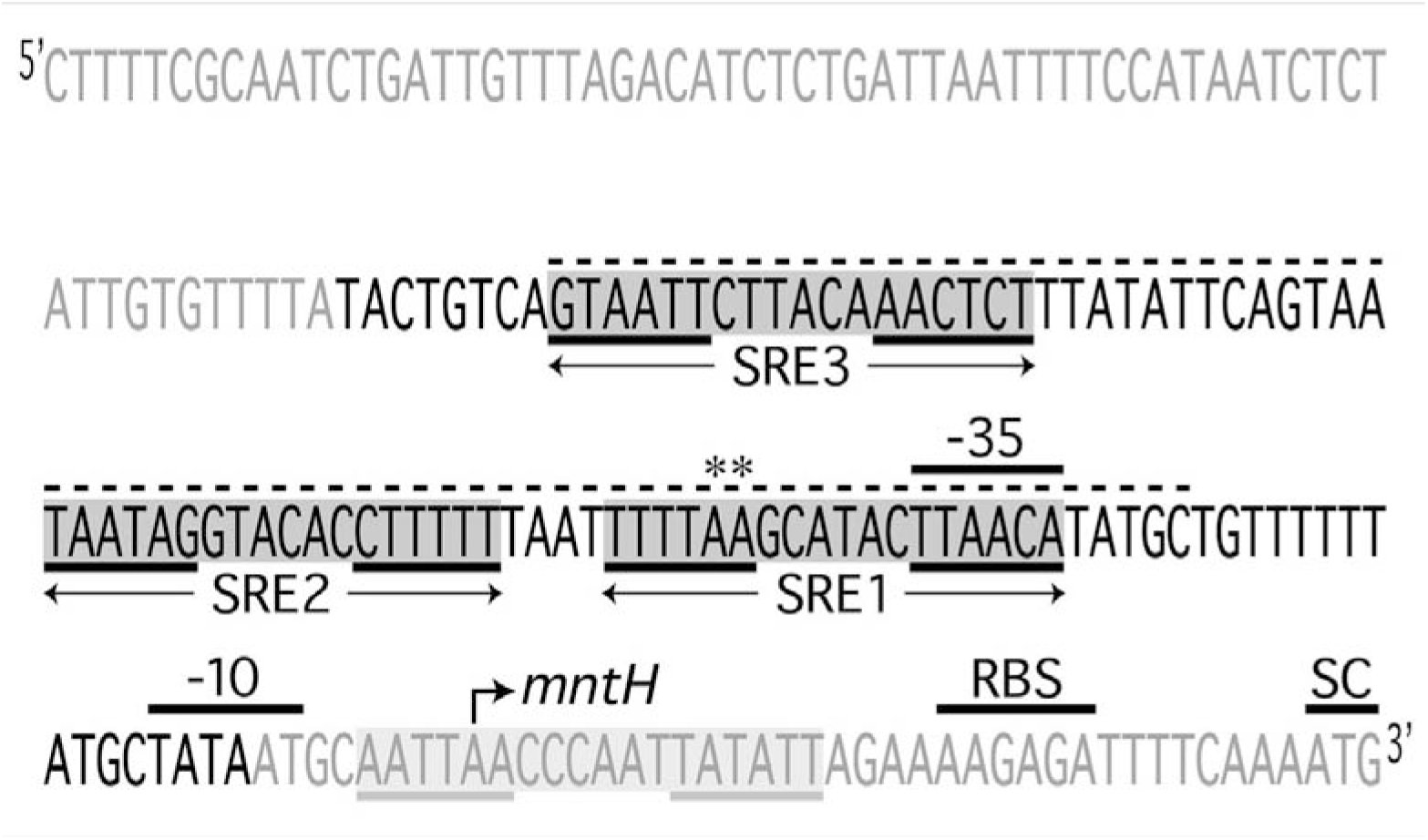
Organization of the *S. mutans mntH* promoter region. Shown are three SREs in the *mntH* promoter region that were predicted *in silico* and subsequently validated in DNA footprinting and gel shift experiments. SRE1 shares overlap with the -35 promoter element. SRE2 is located 4bp adjacent to and upstream of SRE1, and a third SRE is located further upstream of SRE2. The SRE that was predicted by Kajfasz et al (2020) but not validated in this study is located downstream of SRE1 (highlighted in light gray). SREs 1, 2 and 3 share a 6-6-6 binding motif, each composed of different imperfect inverted repeats (underlined). The region of DNA that was protected by SloR in DNA footprinting experiments is designated by the dashed line. The -35 and -10 promoter elements were predicted in 5’ RACE experiments in accordance with the identification of a transcription start site (bent arrow) which defines a 32-bp 5’ untranslated region. Also shown is the predicted ribosome binding site (RBS). Nucleotides designated in black comprise a 100bp target probe that harbors all three SREs, and that was used in gel mobility shift assays. The asterisks indicate the two adenine residues that, when mutated, abrogate SloR binding in EMSA experiments. These same mutations were introduced into the *S. mutans* chromosome (generating GMS2027) which de-repressed *mntH* transcription.

### Regulation of the *S. mutans mntH* gene involves direct SloR binding to at least three SREs in the *mntH* promoter region

A previous report by Kajfasz et al (2020) (7) described direct SloR binding to a 206-bp target probe that harbors the *S. mutans mntH* promoter sequence. To confirm this observation, we performed an EMSA experiment with the same 206-bp target probe, the results of which reveal a robust band shift (Fig. 4) when as little as 60nM SloR was added to the reaction mixture. This band shift was abrogated by the addition of EDTA, consistent with a SloR-DNA interaction that is metal ion-dependent. Having demonstrated that SloR directly binds within the region of the *mntH* promoter, we performed biolayer interferometry to confirm this binding and to define a Kd value for the SloR-*mntH* interaction (Supplementary Fig. 1). Interestingly, the Kd for SloR binding to a 90-bp probe of the *mntH* promoter is ∼33nM, which is similar to that for SloR binding to the 72-bp *sloABC* promoter (∼31nM) which was revealed in a parallel biolayer interferometry assay. Previous Kd determinations for SloR-*sloABC* binding derive from fluorescence anisotropy studies that similarly revealed a strong binding interaction (∼30nM) (17).

**FIG 4.**
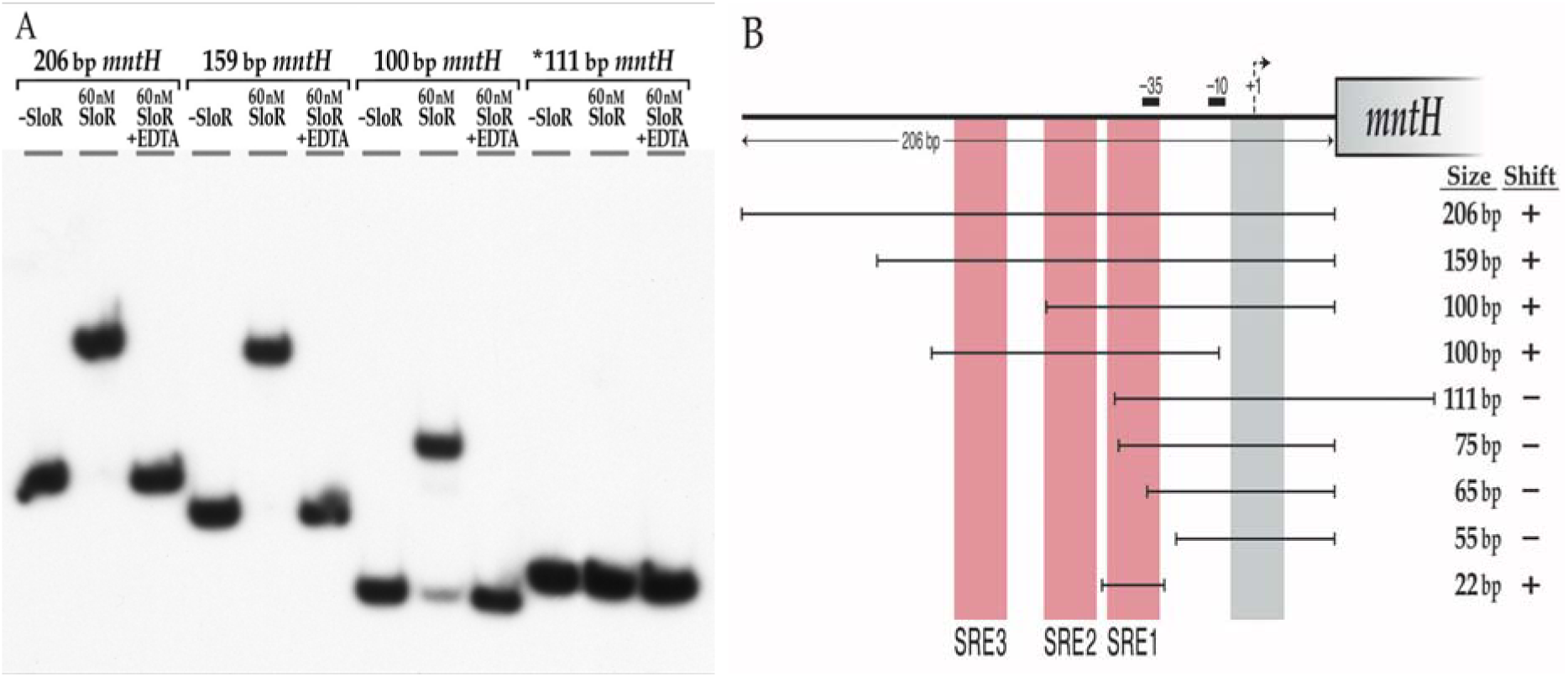
**The *S. mutans* SloR protein binds to the *mntH* promoter fragment**. (a) Amplicons that extend 206-bp, 159-bp, and 100-bp upstream of the *mntH* coding sequence are shifted by SloR. Abrogation of the band shifts by EDTA supports the SloR-DNA interaction as metal ion-dependent. Interestingly, a 111-bp probe that extends 75-bp upstream of the *mntH* start codon and 36-bp downstream of the *mntH* coding sequence was not shifted by SloR (see *). (b) Shown are serial deletion fragments of the 206-bp *mntH* locus that were used as probes in EMSA experiments to reveal the region of SloR binding within the *S. mutans mntH* promoter region. The vertical shading in red indicates the positioning of SREs 1, 2, and 3 relative to the predicted -10 and -35 promoter elements. Shown by the vertical gray bar is a predicted SRE to which SloR does not bind. The (+) designation indicates robust shifting of the probe by 60nM SloR in EMSA experiments; the (–) sign indicates the absence of a shift. Robust band shifts were generated with probes harboring SRE1 but not with probes that excluded this SRE or any part of this binding motif.

Additional EMSA experiments with serial deletion promoter fragments as target probes allowed us to hone in on the specific region(s) in the *mntH* promoter region to which SloR binds (Fig. 4). Via visual inspection, Kajfasz et al. identified a palindrome in the *mntH* promoter region immediately downstream of the predicted -10 promoter element that resembles SREs A, B, and C that precede the *sloABC* genes on the UA159 chromosome (17). We were surprised by the results of EMSA experiments that do not support SloR binding to this sequence (Fig. 4). Instead, we observed robust band shifts with target probes that excluded this predicted SloR binding sequence, and noted the absence of a band shift with probes that harbor only this predicted sequence (data not shown).

Interestingly, target probes that extend less than 75 bp upstream of the *mntH* translation start site failed to generate a band shift in these experiments, whereas those with more than 75 bp of upstream sequence generated a successful shift (Fig. 4a). We therefore focused our SRE search on a ∼25-bp gap region where we identified a putative 6-6-6 binding motif with imperfect inverted repeats (<u>TTTTAA</u>GCATAC<u>TTAACA)</u>. This motif, hereafter called SRE1, shares overlap with the *mntH* -35 promoter element. In fact, we observed robust band shifts when 60nM SloR was added to reaction mixtures containing target probes harboring SRE1, but not with probes that excluded SRE1 or any part thereof (Fig. 4b). To confirm SloR-SRE1 binding, we tested a 22-bp target probe that harbors the full 18-bp SRE1 sequence flanked by two additional base pairs on the 5’ and 3’ ends. Indeed, the 22-bp target probe was successfully shifted in these experiments with as little as 60nM SloR (Fig. 4 and 8). To identify the specific base pair contacts that are required for SloR-SRE1 binding, we performed EMSA experiments with SRE1 and several SRE1 mutant variants as target probes. Specifically, we tested an SRE1 derivative that harbors mutations in the left-most inverted repeat (IR) (<u>TTTTcc</u>GCATAC<u>TTAACA</u>) or in the right-most inverted repeat (<u>TTTTAA</u>GCATAC<u>ggccCA</u>) (data not shown). Both of these IR mutations abolished the band shift, and speak to the importance of SRE 1, and more specifically, to adenine nucleotides for SloR binding.

The results of DNA footprinting experiments support an extended region of protected DNA in the *mntH* promoter that spans 76 base pairs and that can accommodate the binding of at least three SloR homodimers. The footprint includes SRE1 that was shifted by SloR in EMSA studies as well as a second SloR binding sequence, likely <u>TAATAG</u>GTACA<u>CTTTTT</u>, located 4 bp upstream of SRE1 (Fig.5). We went on to investigate whether SloR binding to this upstream sequence, called SRE2, was at all dependent on SloR-SRE1 binding in EMSA experiments with promoter deletion target probes. We observed two distinct band shifts when SloR was added to a reaction mixture containing a wild-type probe with resident SRE1 and SRE2 sequences, consistent with the presence of more than a single SloR binding site in the *mntH* promoter region (Fig. 6). ImageJ analysis revealed a relative band intensity ratio consistent with more prevalent SloR binding to both SREs 1 and 2 as opposed to SRE 1 alone (1.3 to 0.03 AU respectively). A reaction mixture containing 60nM SloR and a target probe with a mutation in SRE1 (but wild-type SRE2) revealed a gel shift pattern consistent with continued SloR binding to more than a single SRE, suggesting the presence of yet another SloR binding site upstream of SRE2 that we named SRE3. A probe with a mutation in SRE2 (but wild-type SRE1) again revealed two band shifts consistent with SloR binding to at least two sites, likely SRE1 and SRE3. Interestingly, ImageJ quantification revealed a high relative intensity band shift that was compromised for SloR binding to SREs 1 and 3 when SRE2 was mutated (2.4 vs.0.7 arbitrary units) as opposed to SloR binding to SREs 1 and 3 when SRE 2 was present in its wild-type configuration (1.3 vs. 0.03 AU). That the upper band shift is compromised when SRE2 is altered supports involvement of SRE2 in SloR binding to SRE3, though to a lesser extent than the role that SRE1 plays in SloR cooperative binding. A target probe with mutations in both SREs 1 and 2 abrogated both band shifts indicating SloR binding to SREs 1 or 2 is a prerequisite for additional SloR-DNA interactions further upstream (ie. SRE3). Taken together, these results support the presence of at least three SREs in the *S. mutans mntH* promoter region, to which SloR appears to be binding cooperatively, favoring binding to SRE1 over SREs 2 and 3.

**FIG 5.**
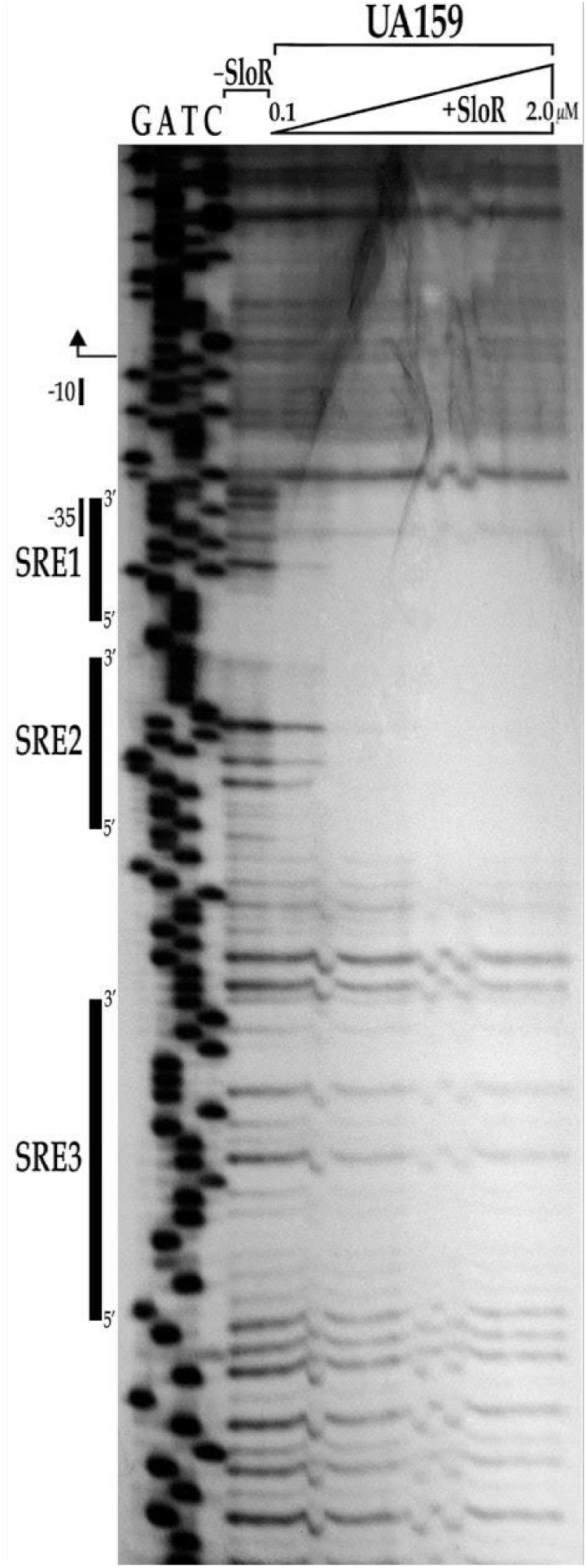
SloR protects a region in the *mntH* promoter harboring at least three SREs. SloR protects a 76-bp region of DNA upstream of the *mntH* coding sequence that includes SREs 1, 2, and 3 as well as the -35 promoter region. A 291-bp amplicon on which the *mntH* promoter is resident was generated with a 5’-end labeled forward primer and a corresponding reverse primer. The radio-labeled amplicon was used in binding reactions with increasing amounts of SloR up to 2 uM. Each reaction along with no protein controls were digested with RQ1 DNase I and resolved on an 8% urea-containing polyacrylamide gel. Shown within the region of protection are at least three footprints, supporting the presence of 3 SREs (indicated by the vertical black bars). The positioning of the -10 and -35 promoter regions are also shown relative to the positioning of the transcription start site (bent arrow).

**FIG 6.**
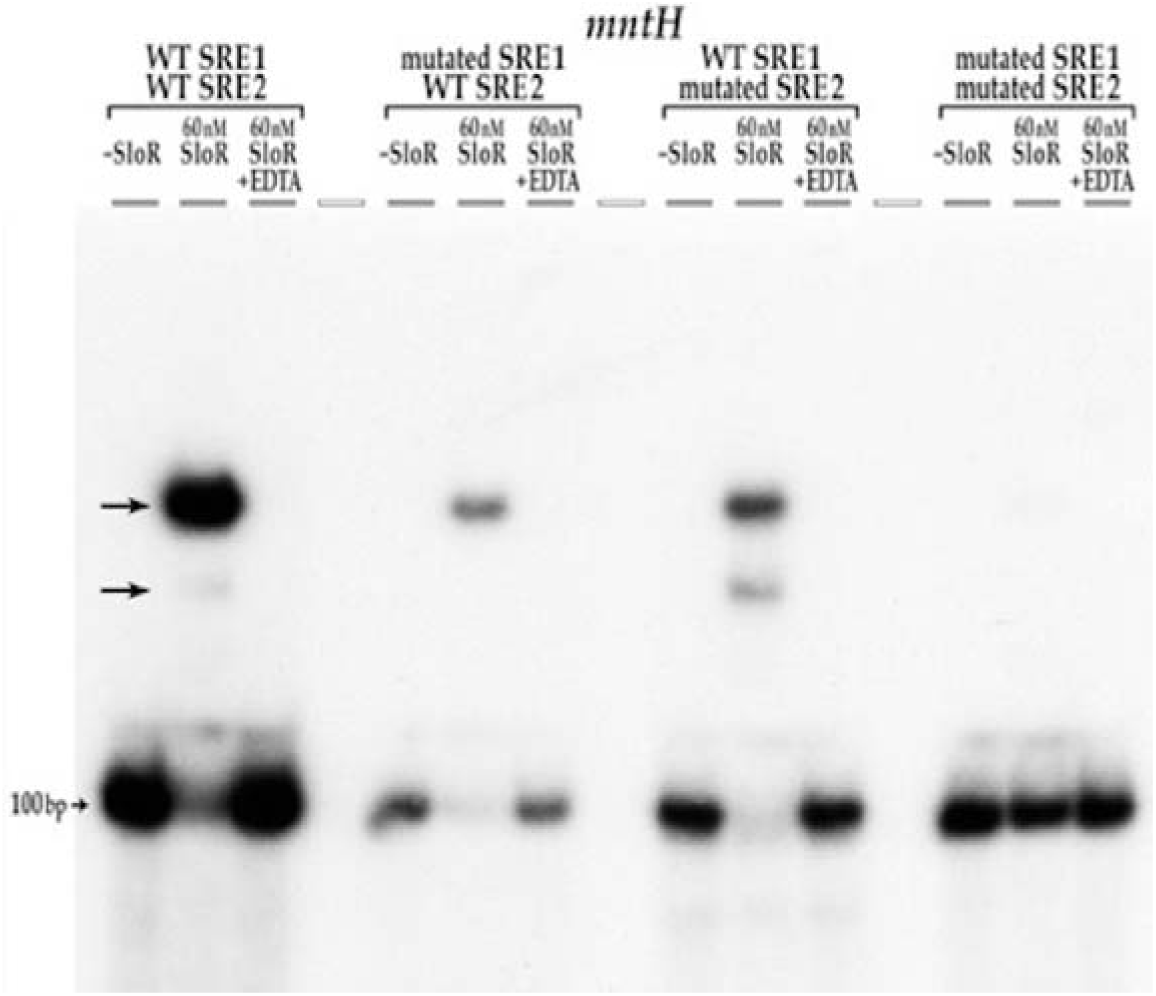
SRE1 mutant variants resident on a 100-bp target probe disrupt cooperative SloR binding to the *mntH* promoter in EMSA experiments. A 100-bp probe that harbors wild-type SREs 1 and 2 produces two visible band shifts (arrows) when 60nM SloR is added to the reaction mixture. The same 100-bp probe with two A-to-C transversion mutations in SRE1 (<u>TTTTcc</u>GCATAC<u>TTAACA</u>) abrogated the lower of the two band shifts. Maintenance of the upper band suggests persistent binding of more than a single SloR homodimer, consistent with the presence of at least one other SRE in addition to SRE 2. Interestingly, the same 100-bp probe with mutations in SRE2 only (<u>TAATtc</u>GTACAC<u>CTTTTT</u>) returns the band shift pattern to that of wild-type, only with different relative band intensities. This result supports SloR binding to SRE 1 as well as to a third SRE upstream of SRE 2, and suggests the loss of cooperative binding when the SloR-SRE 2 interaction is lost or compromised. Abrogation of both band shifts occurs when mutations are introduced into both SREs 1 and 2, consistent with requisite SloR binding to SREs 1 and 2 before the protein can interact with other SREs further upstream. The addition of EDTA to the reaction mixtures abrogates the band shift, indicating that the observed SloR-DNA interactions are metal ion-dependent.

The Kd values that derive from biolayer interferometry are consistent with cooperative SloR binding at the *mntH* locus and implicate a significantly higher SloR binding affinity (and hence a lower Kd) for the SloR-SRE1 interaction than for that of SloR binding to SRE2 and 3. A 22-bp probe that harbors only SRE1 binds to SloR in these experiments, whereas 22-bp probes that harbor either SRE2 or SRE3 alone do not (Supplementary Fig 2). These biolayer interferometry results corroborate the results of EMSA, demonstrating that only SRE1 can support SloR binding when provided as the sole binding site, *in vitro*.

### Mutations in SRE1 and/or SRE2 de-repress transcription of the *S. mutans mntH* gene *in vivo*

We investigated the impact of wild-type and mutant SRE1 and SRE2 variants on *mntH* transcription in real-time semi-quantitative PCR (qRT-PCR) studies. Specifically, we monitored transcription of the *mntH* gene in the wild-type UA159 strain, and in its mutant derivatives GMS2027, GMS2028, and GMS2029. *S. mutans* GMS2027 harbors two adenine substitutions in the leftmost IR of SRE1 (changed to cytosines), to mimic the test probe that was characterized for SloR binding in EMSA experiments. The qRT-PCR results revealed 4.5-fold derepression of *mntH* transcription in GMS2027 compared to its UA159 progenitor (Fig. 7), indicating the essentiality of adenine residues within SRE1 in SloR binding. Transcription of the *mntH* gene in *S. mutans* GMS2028, which harbors two transversion mutations (AG to TC) in the leftmost IR of SRE2, was depressed nearly 3-fold, underscoring the involvement of SRE2 in SloR binding to the *mntH* promoter region. Finally, *mntH* transcription in the GMS2029 SRE variant which harbors both aforementioned mutations in SRE1 and SRE2 respectively, was derepressed more than 3-fold, consistent with a lack of binding to SRE3 when SloR binding to SREs 1 and 2 is disrupted.

**FIG 7.**
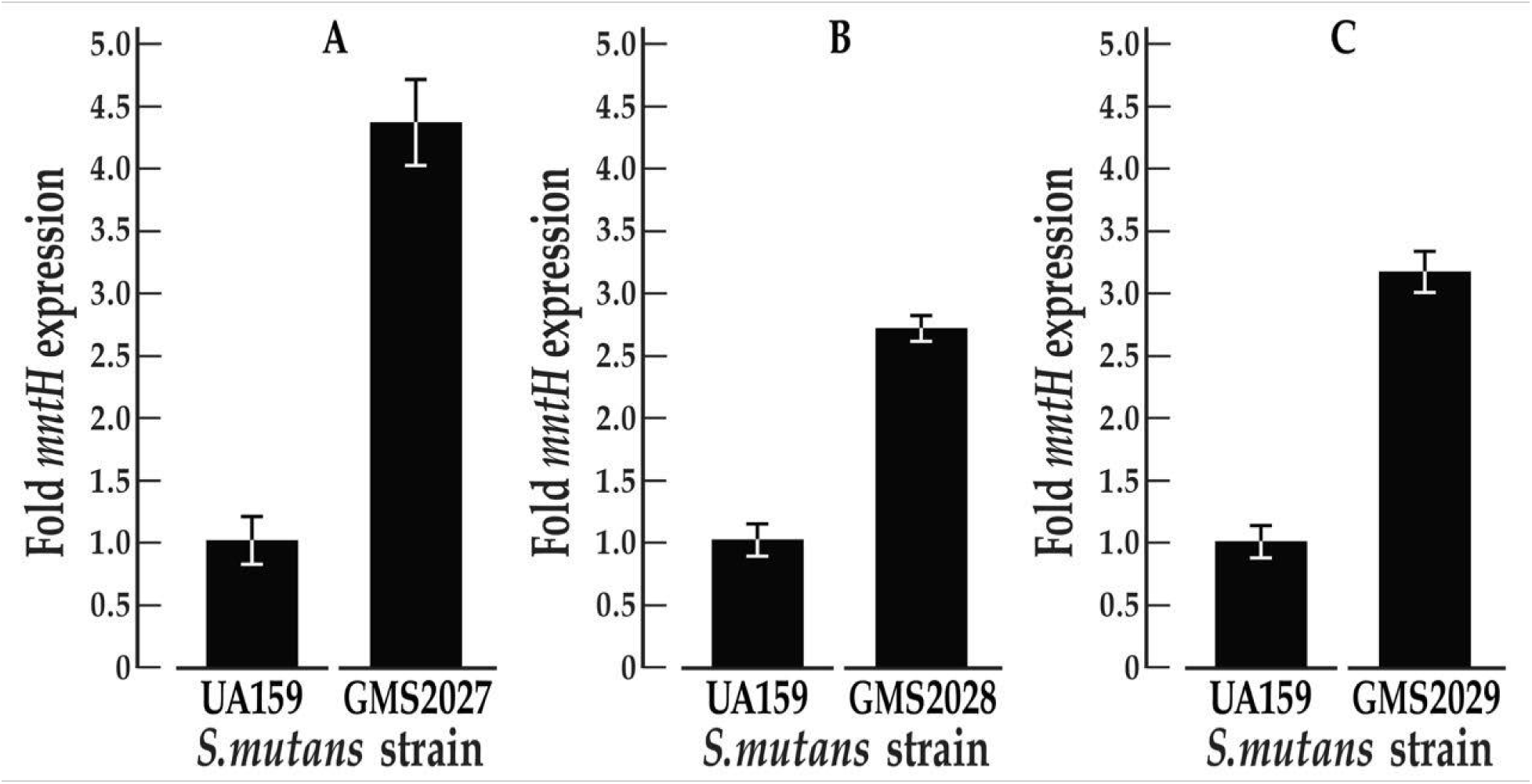
Impact of SRE mutant variants on *S. mutans mntH* transcription *in vivo*. Semi-quantitative real-time PCR experiments were performed in triplicate with the wild-type UA159 *S. mutans* strain and its *mntH* SRE mutant derivatives. The expression of *mntH* was normalized against that of an *hk11* housekeeping gene. (a) The transcription of *mntH* was significantly de-repressed ∼4.5-fold in the GMS2027 mutant strain that harbors an SRE1 mutant variant, compared to wild-type. Transcription of *mntH* in GMS2028 and GMS2029 was also de-repressed compared to wild-type by ∼3-fold. The former harbors two transversion mutations in SRE2 and the latter harbors transversion mutations in both SREs 1 and 2. Shown is the mean *mntH* expression +/- standard deviation that derives from three technical replicates in each of three independent experiments.

### The spacer sequence that separates the inverted repeats (IRs) within an SRE is important but not sufficient for SloR-SRE binding

We conducted EMSA experiments with SRE target probes harboring aberrant IR or spacer sequences to determine whether the SRE spacer in the binding motif is important for SloR-SRE binding. We pursued these experiments because of the degeneracy we observed in the IRs that flank SREs 1, 2 and 3 despite SloR binding to these sequences. Specifically, we constructed a 22-bp target probe with resident SRE1 IR sequences, and a variant 6-bp spacer region. Interestingly, alterations to the SRE1 spacer abolished the robust SloR-SRE1 band shift that we had observed previously with the wild-type SRE1 sequence (Fig. 8). That the SloR-SRE1 band shift was compromised by introducing a unique spacer speaks to the importance of the spacer motif in SloR-SRE binding. We extended these studies to determine whether the SRE1 spacer alone could be sufficient for SloR binding. We did this by generating a target probe composed of IR sequences that we know do not engage SloR with the SRE1 spacer that we know does. When SloR was added to this probe combination, a band shift was still not observed (data not shown). Taken together, these findings support an important role for the SRE spacer in SloR binding but indicate that the spacer alone is not sufficient for the SloR-SRE interaction (Fig. 8).

**FIG 8.**
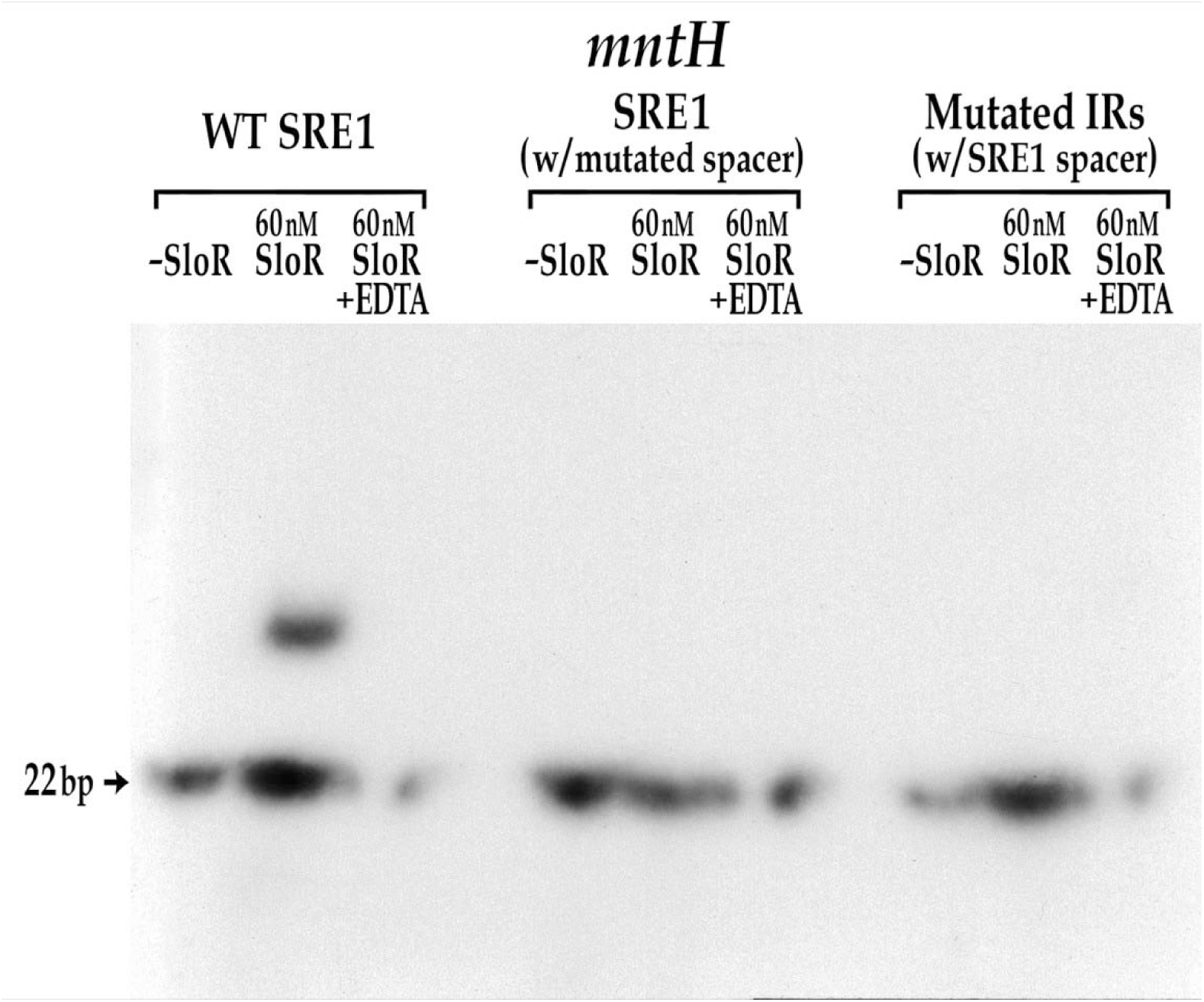
The SRE1 spacer region is important but not sufficient for SloR binding. Wild-type SRE1 (<u>TTTTAA</u>GCATAC<u>TTAACA</u>) in the *mntH* promoter region was successfully shifted by 60nM SloR. In contrast, the same 22-bp probe with a different spacer sequence (<u>TTTTAA</u>CCCAAT<u>TTAACA</u>) was not shifted by SloR. The same 22-bp probe with the SRE1 spacer flanked by unique IR sequences (<u>AATTAA</u>GCATAC<u>TATATT</u>) also failed to generate a band shift. Abrogation of the band shifts by EDTA supports a SloR-DNA interaction that is metal ion-dependent.

## DISCUSSION

Manganese is an essential micronutrient that promotes the survival and pathogenicity of numerous lactic acid bacteria, including *S. mutans* (10). This divalent cation is most widely known as a cofactor for the detoxifying superoxide dismutase (MnSOD) in *S. mutans* that protects the bacterium against oxidative stress (6). Reports in the literature describe SloABC-mediated metal ion transport as the main manganese uptake system in this oral pathogen, although redundant systems have been reported (17). The focus of the present study is on the MntH manganese permease in *S. mutans* which Kajfasz et al. (7) identified as an ancillary transporter of Mn^2+^ via inductively coupled plasma optical emission spectrometry (ICP-OES). In the present study, we performed ^54^Mn uptake experiments with *S. mutans* cells grown in a Mn-replete medium. ^54^Mn accumulation was heightened in the Δ*mntH* single mutant compared to wild-type, and an even greater increase in intracellular ^54^Mn was observed in the double knockout mutant. Taken together, these results support mechanisms for compensatory manganese uptake in *S. mutans* via the SloABC and MntH systems and suggest the existence of possibly other manganese transport systems beyond SloABC and MntH. Compensatory roles for the SloABC and MntH manganese uptake systems are also evident at the level of phenotype. For instance, while we noted a *S. mutans* Δ*mntH*Δ*sloC* double knockout mutant that was significantly compromised for biofilm biomass and architecture when compared with its wild-type UA159 progenitor, its *sloABC* and *mntH* single mutant derivatives were indistinguishable from one another. We propose that the redundancy of manganese uptake systems in *S. mutans* promotes its fitness by allowing it to maintain intracellular Mn^2+^ homeostasis despite the highly transient environment of the human mouth, particularly during periods of feast and famine when Mn^2+^ levels are fluctuating.

Manganese uptake in *S. mutans* occurs primarily via the SloABC system where it is subject to SloR control. Previous work in our laboratory supports SloR binding to so-called SloR recognition elements, or SREs, in the *sloABC* promoter region which represses transcription of the downstream *sloABC* operon (17), presumably via a promoter exclusion mechanism. The results of DNA footprinting assays (17) confirm that SloR binding to the *sloABC* promoter engages each of three inverted hexameric repeat units, each separated by 4bp. Specifically, the three SREs span 72-bp of DNA in the region, and share overlap with the -10 and -35 promoter elements. Each SRE includes an AATTAA IR sequence separated by a 6 bp spacer, comprising a 6-6-6 binding motif. Transcription of the *S. mutans mntH* gene is also subject to SloR control. The results of EMSA experiments support direct SloR binding to a 206-bp *mntH* promoter fragment at protein concentrations as low as 60nM. In fact, the binding studies reported herein support at least three SloR binding sites in the *mntH* promoter within 135-bp of the TSS. SRE1 and its IRs <u>TTTTAA</u> and <u>TTAACA</u> share overlap with the -35 *mntH* promoter element, suggesting that SloR represses *mntH* via steric hindrance (21), a mechanism commonly observed among SloR homologs (22–24). Interestingly, the SRE1 IRs deviate from the IRs of the *sloABC* SRE-B by 4 bp, and the IRs of SREs 2 and 3 deviate from those of the *sloABC* SRE-B by 6 and 11 base pairs, respectively. Consistent with the EMSA findings are the results of biolayer interferometry experiments that reveal SRE1 as the only SRE that can support an independent binding interaction with SloR. In contrast, neither SRE 2 nor 3 alone can bind SloR without what we propose is a stabilizing effect of SloR-SRE1 binding.

The introduction of site-specific mutations into SRE1 abrogated SloR-SRE1 binding but maintained a shifting pattern consistent with SloR binding to more than a single SRE. In contrast, mutations in SRE2 maintained the wild-type gel shift pattern, only with different relative band intensities that speak to a loss of cooperative SloR binding but binding nevertheless to SREs 1 and 3. Mutations in both SREs 1 and 2 abrogated both band shifts. Taken together these findings support the presence of at least three SREs in the *mntH* promoter region and are consistent with cooperative SloR-SRE binding that favors an interaction with SRE1 before engaging with SREs 2 and 3. This is further supported by the biolayer interferometry results.

We reported previously on SloR-SRE binding cooperativity at the *sloABC* promoter (17). Experiments conducted more recently in our laboratory lend even further support to cooperative binding to the promoters that SloR regulates. That is, we demonstrated that C-terminal interactions between adjacent SloR homodimers facilitate SloR-SRE binding and that these interactions are mediated by a quintet of amino acids, including phenylalanine 187 (data not shown). Changing this phenylalanine residue in the C-terminal FeoA domain of the SloR protein with a conservative (F187E) or non-conservative (F187A) amino acid substitution compromised the SloR-SRE interaction in EMSA experiments, and significantly de-repressed *sloABC* transcription in expression profiling studies, consistent with the cooperative binding of SloR homodimers to adjacent SREs (data not shown). Herein we report similar cooperative binding for SloR to at least two adjacent SREs (SREs 1 and 2) in the *mntH* promoter. Cooperative binding at the *S. mutans mntH* locus is further supported by a recent study of MntR, a SloR homolog in *Bacillus subtilis* that binds with cooperativity to the *mneP* promoter (25).

Since SloR binding to SRE1 blocks -35 promoter access to RNA polymerase, we anticipated transcriptional repression of the downstream *mntH* gene in expression profiling experiments. We propose SloR binding to both SRE1 and SRE2 on the *S. mutans* chromosome may fine tune this repression, presumably upon SloR binding to DNA with varying affinities (21). Notably, *in silico* analysis of the *S. mutans* UA159 genome revealed palindromic IR sequences that comprise a putative SRE near the *mntH* -10 promoter element. The IRs of this predicted SRE deviate from the *sloABC* IR sequences by only two base pairs (AATTAA and T<u>TA</u>ATT for *sloABC* versus AATTAA and T<u>AT</u>ATT for *mntH*). Yet, our experimental results do not support SloR binding to this site, despite sharing sequence similarity with the high affinity SRE (Kd = 30nM) at the *sloABC* locus.

The results of biolayer interferometry reveal similar Kd values for SloR binding at the *mntH* and *sloABC* loci. Owing to the sequence divergence between SRE1 and the canonical SRE that precedes the *sloABC* operon, we did not anticipate the near equivalent binding affinities that we observed with SloR. Rather, we anticipated a weaker SloR-SRE1 binding interaction at the *mntH* locus, and hence more relaxed *mntH* transcription that would align with how an ancillary Mn^2+^ permease might fine-tune metal ion import that is otherwise largely determined by SloR-mediated gene control. Instead, the experimental findings reveal SloR binding sequences that can vary significantly and still engage SloR with high affinity. Taken together, these findings support a SloR-SRE interaction in the *S. mutans mntH* promoter region that is likely not wholly sequence-dependent.

We were also intrigued by the evidence that supports an integral role for the SRE spacer in SloR binding. Specifically, we were surprised by the relatively robust binding that we observed for SloR and SRE1 despite the degeneracy of the interrupted palindrome. This led us to hypothesize that the IRs alone may not be fully accountable for SloR-SRE binding and that the 6-bp spacer region may be involved. We performed a spacer mutation experiment to address this hypothesis by substituting the SRE1 spacer with a random sequence of nucleotides (Fig. 8). This spacer alteration effectively abolished SloR binding, demonstrating that the spacer region contributes to SloR binding, and that perfect consensus IRs alone cannot fully explain the SloR-SRE interaction.

Based on what is known about other homodimeric proteins belonging to the DtxR family of metalloregulators, SloR engages the N-terminal helix-turn-helix (H-T-H) motif of each of its monomeric subunits with the major groove of DNA (21). Given that the length of the SRE is 18 bp and that 10.4-bp are required for a single turn of B-DNA, we deduced that the SRE IRs would align with the major groove of the DNA and that the spacer region would align with the minor groove. Although the spacer does not make direct contact with SloR, growing evidence that supports spacer involvement in SloR-DNA binding led us to consider the shape of the DNA and the span of the SloR homodimer in modeling the SloR-SRE binding interaction. Accordingly, we propose that the spacer sequence influences DNA conformation and hence, by inference, the location and accessibility of the SRE IRs to the SloR protein. It is also possible, however, that SloR binding via direct base pair contacts in the major groove (known as direct readout) is less important than previously thought, and that SloR instead interacts with the DNA backbone. Hence, how the spacer impacts the DNA backbone could influence SloR binding affinity as much as the IR sequence (indirect readout). Indirect readout involving SloR recognition of the DNA backbone is reminiscent of the way IdeR in *Mycobacterium* interacts with its DNA target sequence. This IdeR-DNA binding mechanism has its basis in sequence-dependent DNA backbone recognition rather than via base pair contacts with the DNA (26)

In summary, the results of the present study support at least two redundant pathways for manganese import in *S. mutans* that are mediated by the SloABC and MntH systems, and that we believe fine tune metal ion uptake and maximize bacterial fitness. The identification of at least three SREs in the *mntH* promoter region with some degeneracy when compared with the canonical SREs that precede the *S. mutans sloABC* promoter, begs the question of whether sequence dependence is the whole story. Accumulating evidence that points to involvement of the SRE spacer in SloR binding can expand our understanding of the SloR-SRE interaction at the *sloABC* and *mntH* promoters, and potentially others across the *S. mutans* genome. An improved understanding of where and how SloR binds and regulates its gene targets is important because it can inform the development of therapeutics that target the SloR-DNA binding interface, with the ultimate goal of alleviating or preventing dental caries.

## ACKNOWLEDGMENTS

We thank Gary Nelson for figure preparation and Michael Freeman and Will Nemeth for their contributions to this research.

This research was supported by the National Institutes of Health (NIH), grant R01DE014711 to G.A.S., the Middlebury College Biology Department, and the Middlebury College Senior Research Supplement Fund We declare that we have no conflicts of interest to report.

