## Supplemental Material for "SloR-SRE binding to the *S. mutans mntH* promoter is cooperative"

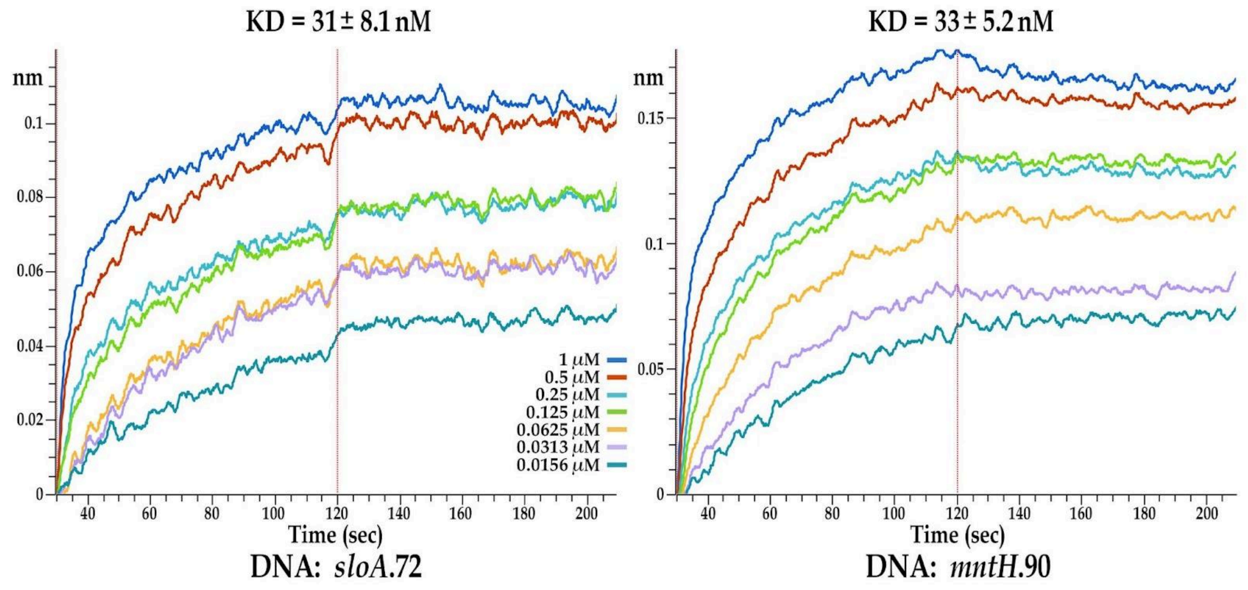

**Supplemental Figure 1.** Binding affinity determinations for the *S. mutans* SloR protein and the *sloABC* and *mntH* promoter probes via biolayer interferometry. Binding measurements were derived from conditions with 5uM SloR protein and a dilution series of DNA target probes ranging from 0.0156uM to 1uM. The real-time binding responses (nM) were recorded over time in seconds (sec) from which the KD value for the interactions was calculated. Shown in (A) is the binding of SloR to a DNA probe spanning 72-bp of the *sloABC* promoter region. Shown in (B) is the binding of SloR to a 90-bp DNA fragment spanning the *mntH* promoter region.

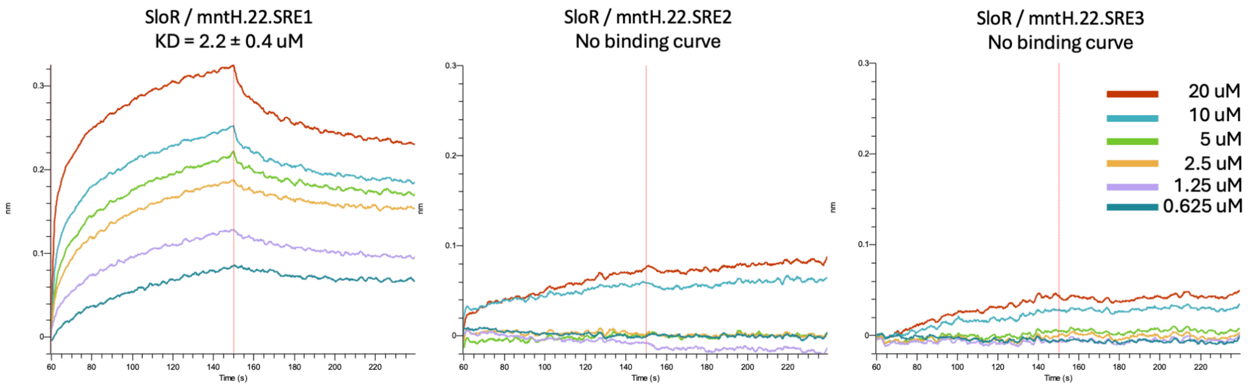

**Supplemental Figure 2.** Binding affinity determinations for the *S. mutans* SloR protein and 22-bp probes in the *mntH* promoter region harboring SRE1, 2, or 3, respectively. Binding measurements were derived from a solution containing 5uM SloR protein and a dilution series of DNA target probes ranging from 0.625uM to 20uM. Real-time binding responses (uM) were recorded over time in seconds (sec) from which KD values were calculated for each interaction. Shown in (A) is the binding interaction between SloR and *mntH* SRE1. (B) and (C) reveal no binding curves for SloR-SRE2 or -SRE3, indicating no SloR binding to either of these *mntH* probes.

**Table 1S. Primers and oligonucleotides used in this study.**

| Application | Name | Nucleotide sequence (5' to 3') | Annealing temp (°C) |
|---|---|---|---|
| **Overlap extension PCR** | **ldhF** | CCGAGCAACAATAACACTC | |
| | **ermR** | GAAGCTGTCAGTAGTATACC | |
| | **770upF** | ATTACAGTTGCGCCAATGATACC | |
| | **770upR-ldh** | GAGTGTTATTGTTGCTCGG*CAATCAGAGCGTTCGTGTTCC | |
| | **770dnF-erm** | TATACTACTGACAGCTTC*TAATCAGAGATGTCTAAACAATCAGATTG | |
| | **770dnR** | TATTGGCAAACGAAGAGCAAG | |
| **Mutagenesis primers** | **770.SRE1.mut.P2** | CAGCATATGTTAAGTATGCggAAAAAT | |
| | **770.SRE1.mut.P3** | ATTTTTccGCATACTTAACATATGCTG | |
| | **700.SRE2.mut.P2** | CATATGTTAAGTATGCTTAAAAATTAAAAAAGGTGTACagATTATTAC | |
| | **700.SRE2.mut.P3** | GTAATAATctGTACACCTTTTTTAATTTTTAAGCATACTTAACATATG | |
| | **700.SRE1.2.mut.P2** | CATATGTTAAGTATGCggAAAAATTAAAAAAGGTGTACagATTATTAC | |
| | **700.SRE1.2.mut.P3** | GTAATAATctGTACACCTTTTTTAATTTTTccGCATACTTAACATATG | |
| **Real-time semi-qRT-PCR** | **mntH.qRT.F** | GTCTTCACTTATTGCCATGC | |
| | **mntH.qRT.R** | GTTGCCATAAGAGCCAATTC | |
| **Amplicon generation for gel shift** | **mntH.1.R** | CATTTTGAAAATCTCTTTTCTAATATAATTG | 61 |
| | **mntH.159.F** | TCTCTATTGTGTTTTATACTGTCAGT | 61 |
| | **mntH.100.F** | AGGTACACCTTTTTTAATTTTTAAGC | 61 |
| | **mntH.75.R2** | ACCTCGCTGAGTGATTGCTT | 68 |
| | **mntH.75.F** | GCATACTTAACATATGCTGTTTTTTATGCT | 68 |
| | **mntH.1.F** | CTTTTCGCAATCTGATTGTTTAG | 61 |
| **Probes in EMSAs** (top & bottom strand) | **mntH.100.WT** | TACTGTCAGTAATTCTTACAAACTCTTTATATTCAGTAATAATAGGTACA CCTTTTTTAATTTTTAAGCATACTTAACATATGCTGTTTTTTATGCTATA<br><br>TATAGCATAAAAAACAGCATATGTTAAGTATGCTTAAAAATTAAAAAAG GTGTACCTATTATTACTGAATATAAAGAGTTTGTAAGAATTACTGACAGT A | |
| | **mntH.100.CC** | TACTGTCAGTAATTCTTACAAACTCTTTATATTCAGTAATAATAGGTACA CCTTTTTTAATTTTTccGCATACTTAACATATGCTGTTTTTTATGCTATA<br><br>TATAGCATAAAAAACAGCATATGTTAAGTATGCggAAAAATTAAAAAAG GTGTACCTATTATTACTGAATATAAAGAGTTTGTAAGAATTACTGACAGT | |

| | | | |
|---|---|---|---|
| | | A | |
| | **mntH.100.CT** | TACTGTCAGTAATTCTTACAAACTCTTTATATTCAGTAATAATctGTACACCTTTTTTAATTTTTAAGCATACTTAACATATGCTGTTTTTTATGCTATA<br><br>TATAGCATAAAAAACAGCATATGTTAAGTATGCTTAAAAATTAAAAAAGGTGTACagATTATTACTGAATATAAAGAGTTTGTAAGAATTACTGACAGTA | |
| | **mntH.100.CC.CT** | TACTGTCAGTAATTCTTACAAACTCTTTATATTCAGTAATAATctGTACACCTTTTTTAATTTTTccGCATACTTAACATATGCTGTTTTTTATGCTATA<br><br>TATAGCATAAAAAACAGCATATGTTAAGTATGCggAAAAATTAAAAAAGGTGTACagATTATTACTGAATATAAAGAGTTTGTAAGAATTACTGACAGTA | |
| | **mntH.22.SRE1.WT** | ATTTTTAAGCATACTTAACATA<br><br>TATGTTAAGTATGCTTAAAAAT | |
| | **mntH.22.SRE1.CC** | ATTTTTccGCATACTTAACATA<br><br>TATGTTAAGTATGCggAAAAAT | |
| | **mntH.22.SRE1.w10.spcr** | ATTTTTAACCCAATTTAACATA<br><br>TATGTTAAATTGGGTTAAAAAT | |
| | **mntH.22.SRE1.10IR** | GCAATTAAGCATACTATATTAG<br><br>CTAATATAGTATGCTTAATTGC | |
| **Query sequence from Kajfasz et al. (2020)** | **206-bp sequence upstream of mntH start codon.** | CTTTTCGCAATCTGATTGTTTAGACATCTCTGATTAATTTTCCATAATCTCTATTGTGTTTTATACTGTCAGTAATTCTTACAAACTCTTTATATTCAGTAATAATAGGTACACCTTTTTTAATTTTTAAGCATACTTAACATATGCTGTTTTTTATGCTATAATGCAATTAACCCAATTATATTAGAAAAGAGATTTTCAAAATG | NA |
| **Primers for 5'RACE** | **770.GSP1.2** | TCAACCTACTGTTAGTAGC | |
| | **770.GSP2.2** | TGTAAGGCAATCCCTGAGCC | |
| | **770_GSP3_nested.2** | TGCCCAGTTTACCAGCCATT | |
| **Primers for DNase I footprinting** | **mntH.FP.F1** | GATTGTTTAGACATCTCTGATTA | |
| | **mntH.FP.R1** | CCTAAAAAAGCTCTTAAAGTTG | |
| **Probes for Biolayer Interferometry** | **mntH.22.SRE1.WT** | ATTTTTAAGCATACTTAACATA | |
| | **mntH.22.SRE2.WT** | AATAATAGGTACACCTTTTTTA | |
| | **mntH.22.SRE3.WT** | CAGTAATTCTTACAAACTCTTT | |
| | **mntH.90.WT** | GTAATTCTTACAAACTCTTTATATTCAGTAATAATAGGTACACCTTTTTTAATTTTTAAGCATACTTAACATATGCTGTTTTTTATGCTA | |

| | | |
|---|---|---|
| | **sloA.72.WT** | AGCCTTAATTAATGTAGATTATATTTTTAATTGAACTGAATTAAAAATATAATCCAATATAATGAATATTTT |
